# Cyclin A/Cdk1 promotes chromosome alignment and timely mitotic progression

**DOI:** 10.1101/2023.12.21.572788

**Authors:** Sarah Y. Valles, Kristina M. Godek, Duane A. Compton

## Abstract

To ensure genomic fidelity a series of spatially and temporally coordinated events are executed during prometaphase of mitosis, including bipolar spindle formation, chromosome attachment to spindle microtubules at kinetochores, the correction of erroneous kinetochore-microtubule (k-MT) attachments, and chromosome congression to the spindle equator. Cyclin A/Cdk1 kinase plays a key role in destabilizing k-MT attachments during prometaphase to promote correction of erroneous k-MT attachments. However, it is unknown if Cyclin A/Cdk1 kinase regulates other events during prometaphase. Here, we investigate additional roles of Cyclin A/Cdk1 in prometaphase by using an siRNA knockdown strategy to deplete endogenous Cyclin A from human cells. We find that depleting Cyclin A significantly extends mitotic duration, specifically prometaphase, because chromosome alignment is delayed. Unaligned chromosomes display erroneous monotelic, syntelic, or lateral k-MT attachments suggesting that bioriented k-MT attachment formation is delayed in the absence of Cyclin A. Mechanistically, chromosome alignment is likely impaired because the localization of the kinetochore proteins BUB1 kinase, KNL1, and MPS1 kinase are reduced in Cyclin A-depleted cells. Moreover, we find that Cyclin A promotes BUB1 kinetochore localization independently of its role in destabilizing k-MT attachments. Thus, Cyclin A/Cdk1 facilitates chromosome alignment during prometaphase to support timely mitotic progression.

## INTRODUCTION

During mitosis a series of spatially and temporally coordinated events occur to ensure that cell division results in two genetically equivalent daughter cells. These events are detectable as morphological changes that have historically been used as criteria to define distinct phases of mitosis. Mitosis begins with prophase, which is defined by the condensation of discernible chromosomes within the nucleus. Prometaphase initiates when the nuclear envelope breaks down and consists of multiple events including bipolar spindle formation, chromosome attachment to spindle microtubules (MTs) at the kinetochore protein complex, chromosome congression to the spindle equator, and the correction of erroneous kinetochore-MT (k-MT) attachments. Metaphase occurs upon completion of chromosome congression and is followed by the segregation of chromosomes to daughter cells in anaphase (Walczak *et al*., 2010). Critically, all these events must occur within an optimal time window to prevent genome instability caused by an accelerated or prolonged mitotic duration (Meraldi *et al*., 2004; Uetake and Sluder, 2010; Daum *et al*., 2011; Sapkota *et al*., 2018; Meitinger *et al*., 2022).

Importantly, the sequence of morphological changes necessary for faithful chromosome segregation are driven by biochemical processes and evidence indicates that each mitotic phase represents a distinct biochemical state within the cell. For example, mitotic entry and the initial phases of mitosis are regulated, in part, by cyclin dependent kinase 1 (Cdk1) activity in complex with Cyclin A or Cyclin B. Because Cyclin A and Cyclin B differ in their temporally controlled proteolytic degradation via the Anaphase Promoting Complex/Cyclosome (APC/C) E3 ubiquitin ligase and 26S proteosome (Clute and Pines, 1999; den Elzen and Pines, 2001; Di Fiore and Pines, 2010), prometaphase is biochemically defined through activity of both Cyclin A/Cdk1 and Cyclin B/Cdk1 while metaphase is defined through predominantly Cyclin B/Cdk1 activity. Cyclin B/Cdk1 activity endures through prometaphase as chromosomes form attachments to the spindle because unattached kinetochores activate spindle assembly checkpoint (SAC) signaling. SAC proteins bind to the APC/C, inhibiting the degradation of Cyclin B/Cdk1 until the SAC is turned off in metaphase when chromosomes are attached to MTs with proper bioriented attachments and congressed to the spindle equator. Once Cyclin B/Cdk1 levels fall below a minimal threshold anaphase onset occurs (Sudakin *et al*., 2001; Musacchio, 2011; London and Biggins, 2014; Lara-Gonzalez *et al*., 2021; McAinsh and Kops, 2023).

In contrast, Cyclin A levels and Cyclin A/Cdk1 activity are high upon entry into mitosis, but Cyclin A destruction begins coincident with mitotic entry. Cyclin A/Cdk1 outcompetes SAC proteins to bind the APC/C. Thus, Cyclin A destruction is not dependent on chromosome biorientation and SAC silencing, rather, it is destroyed in a time dependent manner (den Elzen and Pines, 2001; Wolthuis *et al*., 2008; Di Fiore and Pines, 2010; Zhang *et al*., 2019b; Hégarat *et al*., 2020). Nevertheless, the level and activity of Cyclin A/Cdk1 are important in early mitosis, with known roles in promoting nuclear envelope breakdown and chromatin condensation (Gong and Ferrell, 2010) and regulating k-MT attachment stability (Kabeche and Compton, 2013; Zhang *et al*., 2017). Further, a phosphoproteomic screen identified numerous mitotic proteins that were selectively phosphorylated by Cyclin A/Cdk1 in prometaphase raising the possibility that it has additional functions (Dumitru *et al*., 2017).

Here we investigate if there are other roles of Cyclin A/Cdk1 in mitosis. Using live-cell timelapse imaging and quantitative microscopy of mitotic cancer cells, we determine that Cyclin A/Cdk1 supports timely mitotic progression by promoting chromosome congression and bioriented k-MT attachment formation in prometaphase through the localization of kinetochore components.

## RESULTS

### Cyclin A promotes chromosome congression and mitotic progression

To study the role of Cyclin A/Cdk1 in mitosis, we utilized siRNA to deplete Cyclin A in two cancer cell lines: U2OS and HeLa. We conducted experiments in both cell lines to ensure reproducibility, because non-transformed cells, such as RPE1, did not efficiently enter mitosis following Cyclin A depletion (Hégarat *et al*., 2020). Quantification of Cyclin A levels in immunoblots from asynchronous cells transfected with Cyclin A-targeting siRNA showed a ∼97-98% reduction of Cyclin A (Supplemental Figure 1A). Importantly, knocking down Cyclin A did not cause detectable depletion of Cyclin B (Supplemental Figure 1A) in agreement with previous results (Hégarat *et al*., 2020). Because Cyclin A depletion has been shown to delay or prevent mitotic entry (Gong and Ferrell, 2010; Hégarat *et al*., 2020), we tested if Cyclin A-depleted HeLa and U2OS cells continued to proliferate and enter mitosis. By immunoblotting with antibodies specific to histone H3 Ser10 phosphorylation, a marker for mitotic cells, we confirmed that depletion of Cyclin A did not prohibit mitotic entry in these cell lines (Supplemental Figure 1B). Finally, we analyzed Cyclin A abundance in individual prometaphase cells using immunofluorescence microscopy. This assay also showed that Cyclin A was significantly decreased in cells transfected with Cyclin A-targeting siRNA compared to control transfections (Supplemental Figure 1, C-D and H). Thus, these two cell types can enter mitosis following Cyclin A knockdown, and these results exclude the possibility that the mitotic cells we observe arose from a rare subgroup in which the siRNA was ineffective at reducing Cyclin A levels.

After validating our Cyclin A depletion strategy, we sought to investigate the consequences of Cyclin A depletion in HeLa and U2OS cells as they progressed through mitosis. First, we quantified the percent of HeLa and U2OS cells in each phase of mitosis. This assessment showed a significant increase in the proportion of cells in prometaphase (19% and 21% increase in HeLa and U2OS cells, respectively) in cells depleted of Cyclin A, as visually defined by mitotic cells with one or more unaligned chromosomes (Figure 1, A and B). This suggests that prometaphase duration is prolonged in the absence of Cyclin A.

**Figure 1:**
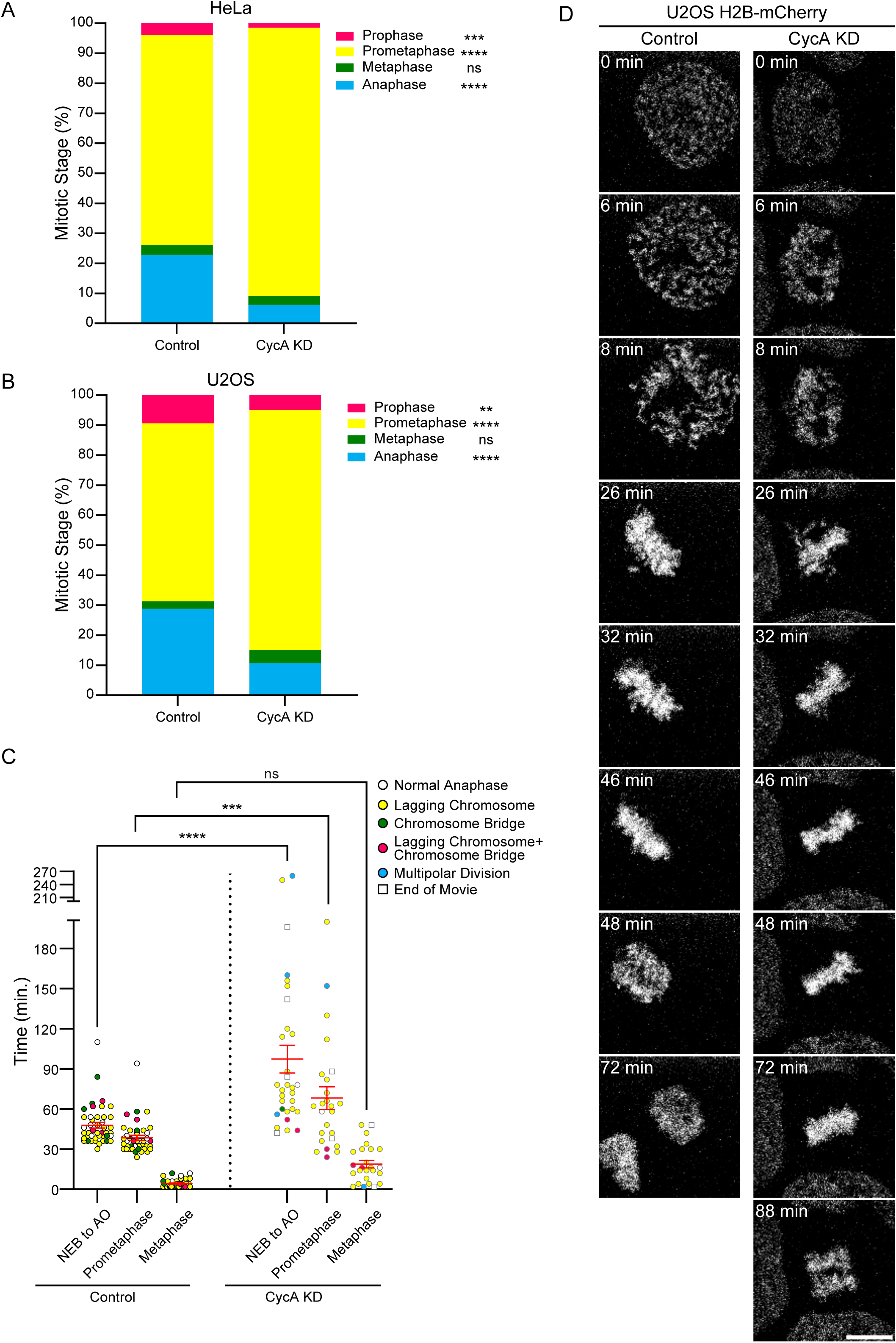
Cyclin A/Cdk1 promotes chromosome congression and mitotic progression. **(A, B)** Percentage of mitotic cells in prophase (magenta), prometaphase (yellow), metaphase (green), and anaphase (blue) in control and Cyclin A depleted (CycA KD) HeLa **(A)** or U2OS cells **(B)**. n > 600 HeLa cells or n > 500 U2OS cells from three independent experiments; ns p > 0.5, **p ≤ 0.01, ***p ≤ 0.001, and ****p ≤ 0.0001 using a two-sided Fisher’s Exact test. **(C)** Total mitotic duration from nuclear envelope breakdown (NEB) to anaphase onset (AO), prometaphase duration, and metaphase duration of control or CycA KD U2OS cells expressing H2B-mCherry from timelapse live-cell fluorescence imaging. Also, anaphases were categorized as normal (white circles) or with an error, including lagging chromosomes (yellow circles), lagging chromosomes and chromosome bridges (magenta circles), or multipolar divisions (blue circles). Cells with anaphases that were not observed before the end of the movie are shown as white squares. For control n = 44 cells from three independent experiments and for CycA KD n = 31 cells from five independent experiments; error bars (red) indicate SEM; ns p > 0.5; ***p ≤ 0.001 and ****p ≤ 0.0001 using a one-way ANOVA and Šídák’s multiple comparisons test. **(D)** Selected panels from timelapse live-cell fluorescence imaging of control or CycA KD U2OS H2B-mCherry cells. Scale bar: 10 µm. See also Supplemental Figure 1.

To corroborate these findings, we utilized live-cell timelapse imaging of asynchronous U2OS H2B-mCherry cells to measure the duration of mitotic events beginning at nuclear envelope breakdown (NEB) and concluding with anaphase onset (AO). These experiments showed a significant elongation in overall mitotic duration (NEB to AO) in cells transfected with Cyclin A siRNA (average = 97.3 minutes) compared to cells transfected with control siRNA (average = 47.8 minutes). This overall delay in mitotic progression was accounted for by a significantly prolonged prometaphase which was distinguished by the presence of unaligned chromosomes (Figure 1, C and D; Supplemental Figure 1E). Consistent with previous studies, this live-cell imaging approach also showed that the cells depleted of Cyclin A displayed elevated rates of lagging chromosomes in anaphase compared to control cells (Kabeche and Compton, 2013; Zhang *et al*., 2017) (Supplemental Figure 1, F and G). In parallel, immunofluorescence analysis on the cells used for live-cell imaging also validated that they lacked detectable Cyclin A in prometaphase (Supplemental Figure 1H).

Chromosome alignment arises through k-MT attachment and biorientation (Kapoor *et al*., 2006; Maiato *et al*., 2017; Craske and Welburn, 2020), therefore, we utilized immunofluorescence analysis of calcium-stable spindle MTs to assess k-MT attachments in mitotic cells (Warren *et al*., 2020). In agreement with previous results, chromosomes aligned at the spindle equator exhibited bioriented k-MT attachments in both control and Cyclin A-depleted HeLa and U2OS cells (Figure 2, A and B) (Kabeche and Compton, 2013). There was also no detectable difference in intercentromere distance on bioriented chromosomes in prometaphase between cells depleted of Cyclin A or control cells, although we detected a small increase in intercentromere distance in metaphase of Cyclin A-depleted cells (Figure 2C). Thus, even in the absence of Cyclin A, the bioriented k-MT attachments that form generate sufficient tension between sister centromeres. On unaligned chromosomes in prometaphase, k-MT attachments exhibited a variety of attachment types, including erroneous monotelic, syntelic, lateral, and unattached configurations in Cyclin A-depleted cells (Figure 2, A-B and D). These results are consistent with previous work demonstrating that Cyclin A-deficient cells exhibit hyper-stable k-MT attachments in prometaphase which erodes the correction of erroneous k-MT attachments (Kabeche and Compton, 2013; Zhang *et al*., 2017). Taken together, the data presented here extends those findings to indicate that Cyclin A, and by extension Cyclin A/Cdk1 activity, promotes chromosome congression and the establishment of bioriented k-MT attachments to support timely progression through prometaphase.

**Figure 2:**
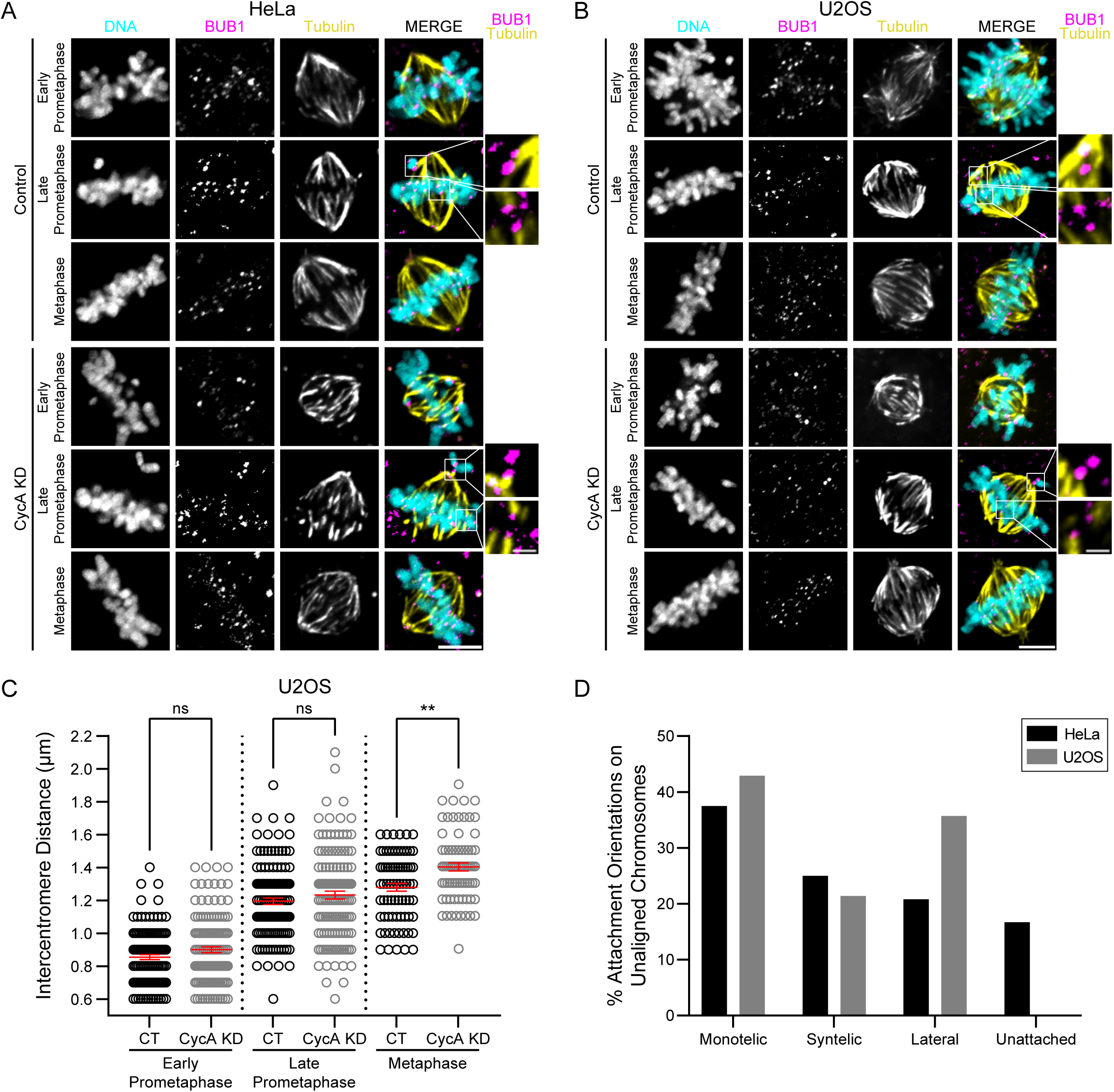
Cyclin A/Cdk1 facilitates the formation of bioriented kinetochore microtubule attachments. **(A, B)** Representative images of microtubule attachments in early prometaphase, late prometaphase (with one unaligned chromosome), and metaphase in control and Cyclin A depleted (CycA KD) HeLa **(A)** or U2OS cells **(B)**. Shown is DNA (cyan), BUB1 (magenta), and microtubules (yellow). For HeLa cells **(A)**, insets show a syntelic k-MT attachment (top) and bioriented k-MT attachment (bottom) in control cells and a monotelic k-MT attachment (top) and a bioriented k-MT attachment (bottom) in CycA KD cells. For U2OS cells **(B)**, insets show monotelic k-MT attachments (top) and bioriented k-MT attachments (bottom) in both the control and CycA KD U2OS cells. Scale bars: 5 µm or 1 µm for insets. **(C)** Quantification of intercentromere distance in early prometaphase, late prometaphase, and metaphase control (black) or CycA KD (gray) U2OS cells. n = 75-130 pairs of centromeres from 15-26 cells per mitotic phase. These measurements were performed on cells from the four independent experiments in Figure 7; multipolar cells were excluded; ns p > 0.05, **p ≤ 0.01, using a one-way ANOVA and Šídák’s multiple comparisons test; error bars (red) indicate SEM. **(D)** Percent of k-MT attachments on unaligned chromosomes classified as monotelic, syntelic, lateral, or unattached in CycA KD HeLa (black) and CycA KD U2OS (gray) cells. HeLa n = 24 unaligned chromosomes from 13 cells and U2OS n = 14 unaligned chromosomes from 6 cells.

### Kinetochore composition relies on Cyclin A

Chromosome congression relies on the functional activity of many proteins, and previous reports show that the kinetochore kinase BUB1 has a role in chromosome congression in addition to a role in spindle assembly checkpoint (SAC) signaling (Johnson *et al*., 2004; Meraldi and Sorger, 2005; Klebig *et al*., 2009; Ricke *et al*., 2012; Baron *et al*., 2016; Nguyen *et al*., 2021). To test if the delay in chromosome congression induced by depletion of Cyclin A involved changes in levels of BUB1 localization at kinetochores, we quantified BUB1 intensity at kinetochores in HeLa and U2OS cells following transfection with Cyclin A siRNA. We found that in early and late prometaphase as well as metaphase depletion of Cyclin A resulted in a significant reduction of BUB1 levels at kinetochores compared to control cells (Figure 3, A-B and E-F). This was particularly evident in early prometaphase where BUB1 levels were decreased by 42% or 44% in Cyclin A-depleted HeLa or U2OS cells, respectively. We also selectively measured BUB1 levels at kinetochores of unaligned chromosomes relative to aligned chromosomes to explore whether the depletion of Cyclin A differentially affected the kinetochore localization of BUB1 dependent on chromosome alignment. For this experiment, we focused on late prometaphase, defined by the presence of a metaphase plate but with one or a few unaligned chromosomes. This analysis showed that BUB1 intensity on kinetochores of the unaligned chromosomes was 19% and 45% lower in HeLa and U2OS cells, respectively, in Cyclin A-depleted cells compared to control cells (Figure 3, C-D and G-H). Thus, Cyclin A depletion reduces BUB1 levels on kinetochores of aligned and unaligned chromosomes. Importantly, the observed decrease in BUB1 at kinetochores was not a consequence of an overall inability to properly form kinetochores because CENP-A localization increased slightly in Cyclin A-depleted HeLa and U2OS cells (Figure 3, A and E; Supplemental Figure 2, A and B). Moreover, the reduction of BUB1 at kinetochores we observed is likely insufficient to abrogate the SAC because near total depletion of BUB1 is required to show a SAC defect (Zhang *et al*., 2019a).

**Figure 3:**
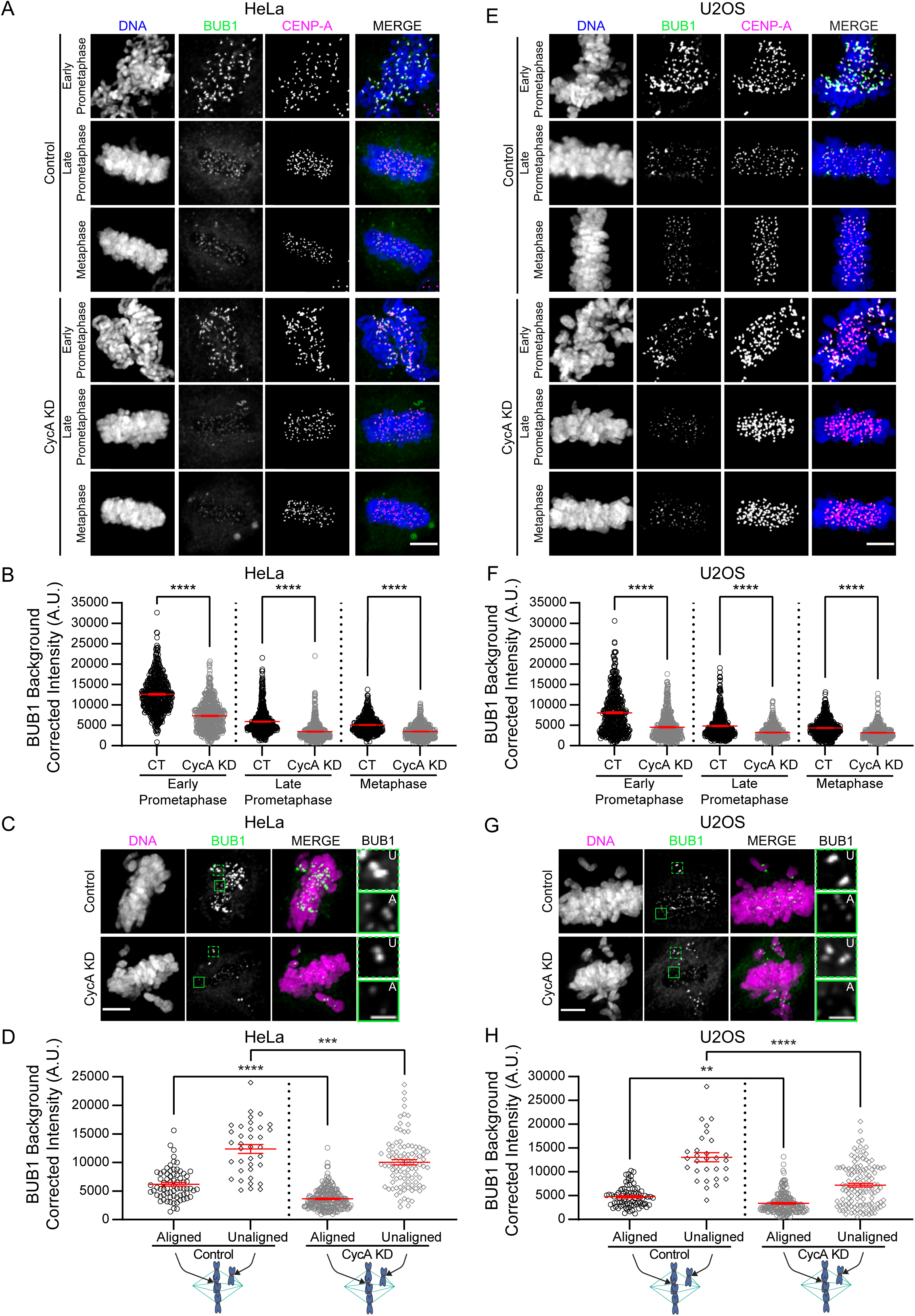
Cyclin A/Cdk1 promotes BUB1 localization to kinetochores. **(A)** Representative images of control or Cyclin A depleted (CycA KD) HeLa cells. Shown is DNA (blue), BUB1 (green), and CENP-A (magenta) in early prometaphase, late prometaphase, and metaphase. **(B)** Quantification of BUB1 protein levels at kinetochores of control (CT, black circles) and CycA KD (gray circles) HeLa cells in early prometaphase, late prometaphase, and metaphase. n > 500 kinetochores from 29-33 cells per mitotic phase. **(C)** Representative images of control and CycA KD HeLa cells with unaligned (U, dark green) and aligned (A, light green) chromosomes in late prometaphase. Shown is DNA (magenta) and BUB1 (green). Insets are magnified views. **(D)** Quantification of BUB1 protein levels at kinetochores from aligned (circles) and unaligned (diamonds) chromosomes in late prometaphase. From 7 control HeLa cells, n = 71 kinetochores from aligned chromosomes and n = 36 kinetochores from unaligned chromosomes. From 22 CycA KD HeLa cells, n = 223 kinetochores from aligned chromosomes and n = 97 kinetochores from unaligned chromosomes. **(E)** Representative images of control or Cyclin A depleted (CycA KD) U2OS cells. Shown is DNA (blue), BUB1 (green), and CENP-A (magenta) in early prometaphase, late prometaphase, and metaphase. **(F)** Quantification of BUB1 protein levels at kinetochores of control (CT, black circles) and CycA KD (gray circles) U2OS cells in early prometaphase, late prometaphase, and metaphase. n = 500 kinetochores from 30-32 cells per mitotic phase. **(G)** Representative images of control and CycA KD U2OS cells with unaligned (U, dark green) and aligned (A, light green) chromosomes in late prometaphase. Shown is DNA (magenta) and BUB1 (green). Insets are magnified views. **(H)** Quantification of BUB1 protein levels at kinetochores from aligned (circles) and unaligned (diamonds) chromosomes in late prometaphase. From 8 control U2OS cells, n = 80 kinetochores from aligned chromosomes and n = 29 kinetochores from unaligned chromosomes. From 18 CycA KD U2OS cells, n = 184 kinetochores from aligned chromosomes and n = 124 kinetochores from unaligned chromosomes. Data from **(B, D, F,** and **H)** are from three independent experiments; **p ≤ 0.01, ***p ≤ 0.001, and ****p ≤ 0.0001 using a one-way ANOVA and Šídák’s multiple comparisons test; error bars (red) indicate SEM. Scale bars **(A, C, E,** and **G)**: 5 µm or 1 µm for insets. See also Supplemental Figure 2.

Next, we tested if the reduction in BUB1 at kinetochores caused by Cyclin A depletion altered the phosphorylation of histone H2A Thr120 (H2ApT120). Phosphorylation of histone H2A T120 by BUB1 is important for the kinetochore-proximal binding of the chromosomal passenger complex (CPC) (Yamagishi *et al*., 2010; Carmena *et al*., 2012; Bekier *et al*., 2015; Baron *et al*., 2016; Hadders *et al*., 2020). We quantified the abundance of H2ApT120 that co-localized with BUB1 at kinetochores and observed a significant reduction in H2A T120 phosphorylation of 60% and 54% in HeLa cells upon Cyclin A depletion in early and late prometaphase, respectively (Figure 4, A and B). The extent of reduction in histone H2A T120 phosphorylation in these cells relative to the reduction of total BUB1 at kinetochores was not strictly proportional suggesting that the decrease in histone H2A T120 phosphorylation results from Cyclin A affecting additional factors (possibly BUB1 kinase activity or an opposing phosphatase) in addition to the reduction of BUB1 abundance at kinetochores (Figure 4C). There was also a significant reduction of histone H2A T120 phosphorylation in U2OS cells in Cyclin A-depleted cells although the extent of this reduction was more modest than in HeLa cells (Figure 4, D and E). Accordingly, the decrease in histone H2A T120 phosphorylation in U2OS cells relative to the reduction of total BUB1 at kinetochores was inverted suggesting that the remaining BUB1 at kinetochores in Cyclin A-depleted U2OS cells is sufficient to maintain some histone H2A T120 phosphorylation (Figure 4F).

**Figure 4:**
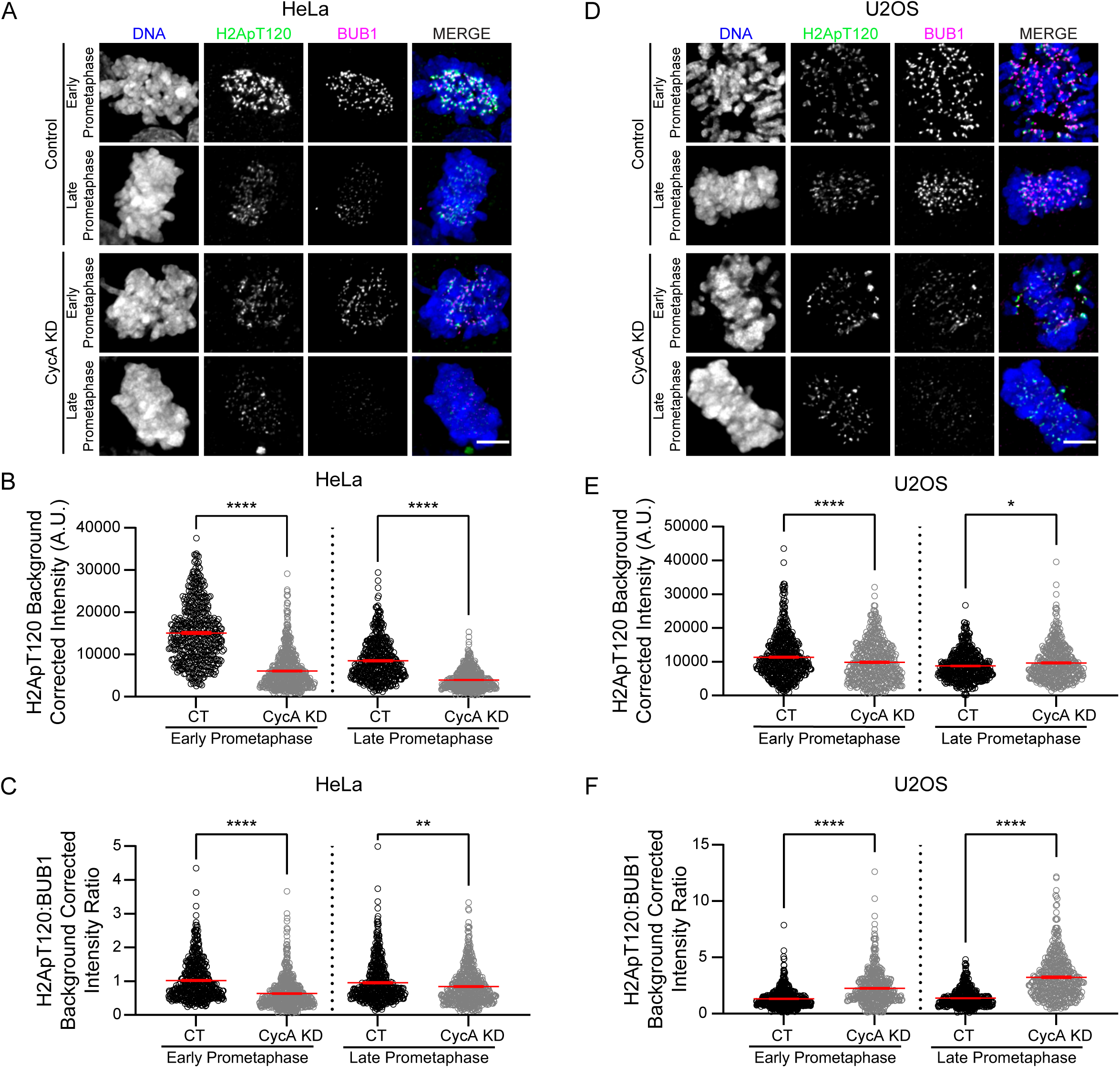
Cyclin A/Cdk1 increases the BUB1 dependent phosphorylation of H2A Thr 120 at kinetochores. **(A)** Representative images of control and Cyclin A depleted (CycA KD) HeLa cells. Shown is DNA (blue), H2A pT120 (green), and BUB1 (magenta) in early and late prometaphase. **(B)** Quantification of H2A phosphorylated on T120 at kinetochores of control (black circles) and CycA KD (gray circles) HeLa cells. n > 500 kinetochores from 30-32 cells per mitotic phase. **(C)** Ratio of H2A pT120 to BUB1 intensity levels at kinetochores of control (black circles) and CycA KD (gray circles) HeLa cells. n >500 kinetochores from 30-32 cells per mitotic phase. **(D)** Representative images of control and Cyclin A depleted (CycA KD) U2OS cells. Shown is DNA (blue), H2A pT120 (green), and BUB1 (magenta) in early and late prometaphase. **(E)** Quantification of H2A phosphorylated on T120 at kinetochores of control (black circles) and CycA KD (gray circles) U2OS cells. n > 450 kinetochores from 27-31 cells per mitotic phase. **(F)** Ratio of H2A pT120 to BUB1 intensity levels at kinetochores of control (black circles) and CycA KD (gray circles) HeLa cells. n >450 kinetochores from 27-31 cells per mitotic phase. Data from **(B and C)** and **(E and F)** are from three independent experiments; *p ≤ 0.05 and ****p ≤ 0.0001 using a one-way ANOVA and Šídák’s multiple comparisons test; error bars (red) indicate SEM. Scale bars **(A** and **D)**: 5 µm.

The canonical localization pathway of BUB1 to kinetochores relies on KNL1. KNL1 is a large scaffolding protein at the kinetochore that is phosphorylated by MPS1 kinase at MELT motifs to provide a binding platform for the BUB1:BUB3 complex (Primorac *et al*., 2013; Vleugel *et al*., 2013; Zhang *et al*., 2013; Ghongane *et al*., 2014; Nijenhuis *et al*., 2014; Fischer, 2023). Using quantitative immunofluorescence analysis, we find that the intensity level of KNL1 at kinetochores is significantly reduced in Cyclin A-depleted early and late prometaphase HeLa and U2OS cells compared to control cells (43 and 48% HeLa and 20 and 27% U2OS, respectively) (Figure 5, A-D). The reduction of KNL1 at kinetochores may also contribute to the delay in chromosome alignment as depletion of KNL1 itself causes chromosome alignment defects (Caldas *et al*., 2013). We also measured the intensity of KNL1 phosphorylation at two MELT sites (Thr 943 and Thr 1155 with identical sequence motifs that are both recognized by the same antibody) at kinetochores in Cyclin A-depleted cells (Nijenhuis *et al*., 2014). We find a significant decrease in MELT phosphorylation (pMELT) in both HeLa and U2OS cells in early prometaphase of Cyclin A-depleted cells compared to control cells (Figure 5, E and G). Moreover, the decrease in pMELT exceeds the reduction in total KNL1 abundance at kinetochores (compare Figure 5, B and D to F and H) suggesting that Cyclin A contributes to KNL1 MELT phosphorylation through multiple pathways during early prometaphase. In contrast, in late prometaphase, pMELT was reduced to levels proportional to the reduction in total KNL1 levels (23% HeLa and 12% U2OS) suggesting that the decrease in phosphorylation of MELT sites at this stage of mitosis is explained by the reduced abundance of KNL1 at kinetochores in Cyclin A depleted cells. These data do not comprehensively assess all MELT sites on KNL1, but the pMELT antibody used in this assay recognizes a MELT repeat that can rescue chromosome alignment defects in the absence of other MELT domains (Vleugel *et al*., 2013). Together, these results show that Cyclin A/Cdk1 activity promotes the localization of KNL1 and BUB1, through phosphorylation of MELT sites on KNL1, in early mitosis.

**Figure 5:**
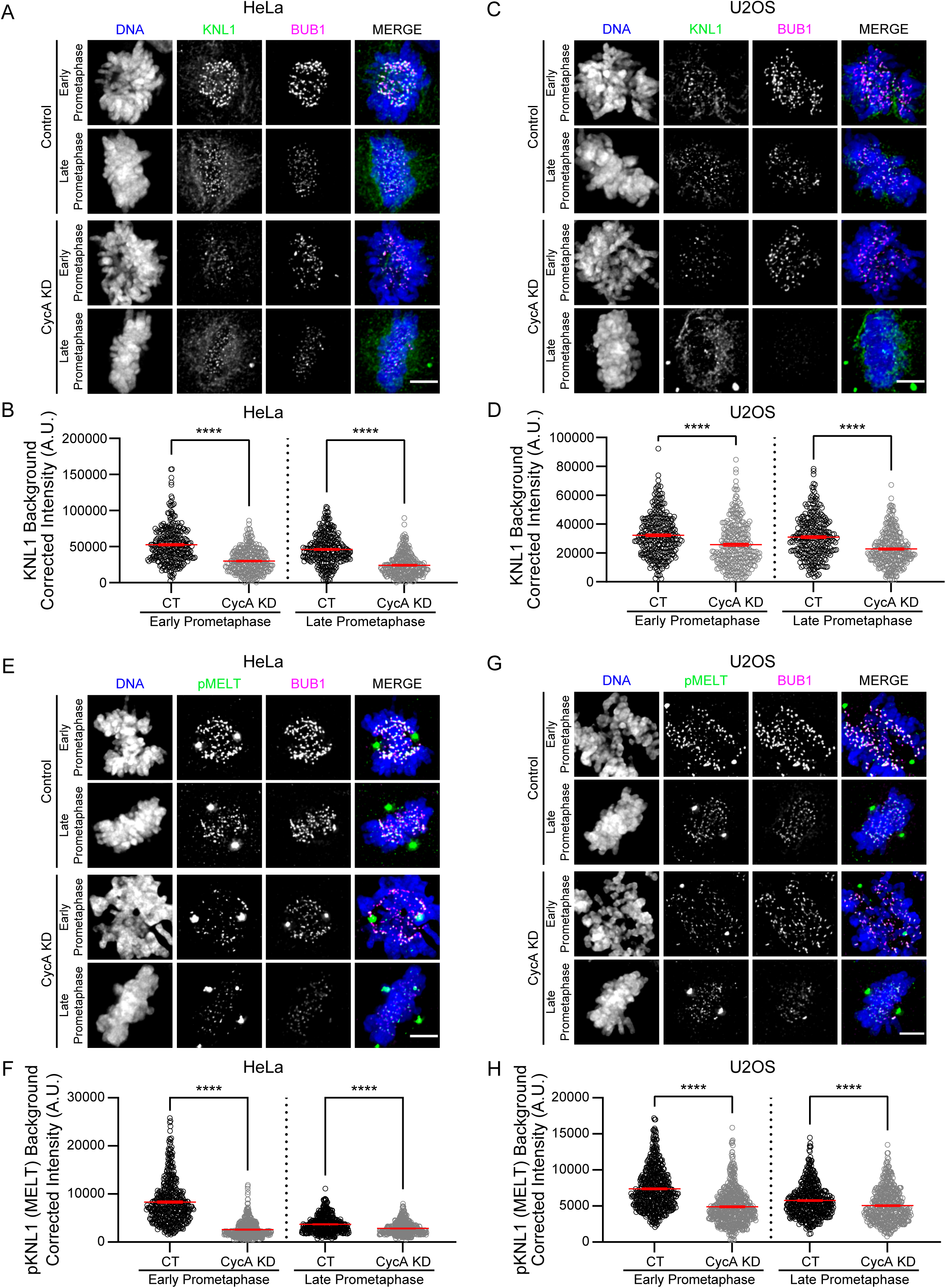
Cyclin A/Cdk1 promotes KNL1 localization to kinetochores and phosphorylation of KNL1 MELT motifs. **(A)** Representative images of control and Cyclin A depleted (CycA KD) HeLa cells. Shown is DNA (blue), KNL1 (green), and BUB1 (magenta) in early and late prometaphase. **(B)** Quantification of total KNL1 protein levels at kinetochores of control (CT, black circles) and CycA KD (gray circles) HeLa cells. n = 300 kinetochores from 30 cells per mitotic phase. **(C)** Representative images of control and CycA KD U2OS cells. Shown is DNA (blue), KNL1 (green), and BUB1 (magenta) in early and late prometaphase. **(D)** Quantification of total KNL1 protein levels at kinetochores of control (CT, black circles) and CycA KD (gray circles) U2OS cells. n = 300 kinetochores from 30 cells per mitotic phase. **(E)** Representative images of control and CycA KD HeLa cells in early and late prometaphase. Shown is DNA (blue), KNL1 pMELT (green), and BUB1 (magenta). **(F)** Quantification of KNL1 pMELT levels at kinetochores of CT (black circles) and CycA KD (gray circles) HeLa cells. n > 450 kinetochores from 30-33 cells per mitotic phase. **(G)** Representative images of control and CycA KD U2OS cells in early and late prometaphase. Shown in DNA (blue), KNL1 pMELT (green), and BUB1 (magenta). **(H)** Quantification of KNL1 pMELT levels at kinetochores of CT (black circles) and CycA KD (gray circles) U2OS cells. n > 450 kinetochores from 31-33 cells per mitotic phase. Data from **(B, D, F,** and **H)** are from three independent experiments; ****p ≤ 0.0001 using a one-way ANOVA and Šídák’s multiple comparisons test; error bars (red) indicate SEM. Scale bars **(A, C, E,** and **G)**: 5 µm.

MPS1 kinase is responsible for the phosphorylation of KNL1 on MELT sites during mitosis (Zhang *et al*., 2013; Vleugel *et al*., 2015). Thus, we used immunofluorescence to test if a decrease in MPS1 kinase localization to kinetochores could be an explanation for the reduction in KNL1 pMELT levels. We focused on early prometaphase because that is when MPS1 displays the most pronounced kinetochore localization. We find a significant decrease in MPS1 intensity at kinetochores in Cyclin A-depleted HeLa cells compared to control cells (Figure 6, A and B). The extent of MPS1 reduction in Cyclin A-depleted cells was less than the reduction of KNL1 pMELT, suggesting that the catalytic activity is reduced more profoundly than the total protein localized and/or the presence of ongoing phosphatase activity may be limiting the extent of KNL1 pMELT at kinetochores in Cyclin A-depleted cells (Kabeche and Compton, 2013; Kucharski *et al*., 2022).

**Figure 6:**
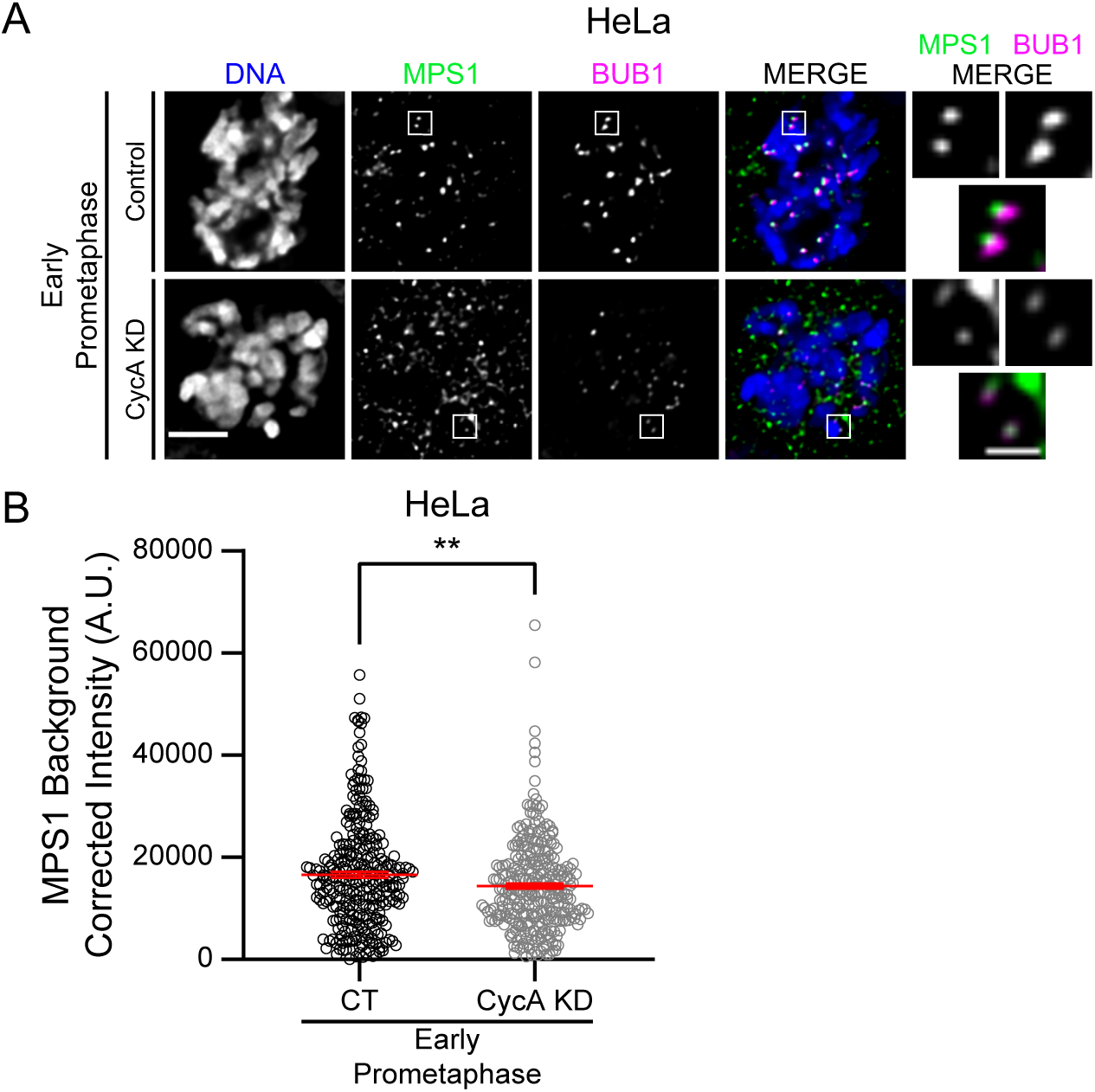
Cyclin A/Cdk1 enhances MPS1 localization to kinetochores. **(A)** Representative images of control and Cyclin A depleted (CycA KD) HeLa cells. Shown is DNA (blue), MPS1 (green), and BUB1 (magenta) in early prometaphase. Scale bars: 5 µm and 1 µm for inset. **(B)** Quantification of MPS1 protein levels at kinetochores of (CT, black circles) and CycA KD (gray circles) HeLa cells. n = 300 kinetochores from 30 cells per condition. Data are from three independent experiments; **p ≤ 0.01 using a two-tailed Student’s t test; error bars (red) indicate SEM.

### Cyclin A-dependent kinetochore localization of BUB1 is independent of k-MT attachment stability

We previously demonstrated that Cyclin A/Cdk1 activity destabilizes k-MT attachments during prometaphase (Kabeche and Compton, 2013). Here, we show that Cyclin A/Cdk1 activity promotes the recruitment of KNL1, BUB1, and MPS1 to kinetochores in prometaphase of both U2OS and HeLa cells (Figures 3, 5, and 6). Thus, Cyclin A/Cdk1 activity could promote the localization of these kinetochore components directly or indirectly as a consequence of altering k-MT attachment stability. To test this, we determined whether BUB1 kinetochore localization is dependent on k-MT attachment stability. We used quantitative immunofluorescence to measure BUB1 intensity under conditions that alter k-MT attachment stability without altering Cyclin A levels. For this purpose, we treated cells with a short-term low-dose of Taxol which causes a similar, approximately 1.8-fold increase, in k-MT attachment stability (Warren *et al*., 2020) as the approximately 1.6-fold increase in k-MT attachment stability observed in Cyclin A-depleted cells in prometaphase (Kabeche and Compton, 2013).

Upon short-term Taxol treatment, BUB1 intensity at kinetochores decreased by only 15% in early prometaphase, did not significantly change in late prometaphase, and increased slightly in metaphase compared to control cells (Figure 7, A and B). Meanwhile, BUB1 intensity at kinetochores in Cyclin A-depleted cells decreased by approximately 40% relative to Taxol-treated cells in all phases of mitosis examined. Thus, BUB1 localization to kinetochores is minimally affected by low dose Taxol treatment compared to Cyclin A depletion, despite the fact that these two treatments cause a similar magnitude in change to the stability of k-MT attachments. These data show that Cyclin A/Cdk1 promotes BUB1 localization to kinetochores through a mechanism that is independent of how it alters the stability of k-MT attachments during prometaphase (Figure 7). Similarly, we find that CENP-A levels were minimally affected by Taxol treatment compared to Cyclin A depletion (Supplemental Figure 3). Also, of note, we observed multipolar spindles in some Taxol-treated and Cyclin A-depleted cells, as well as in some control cells, but all cells analyzed in metaphase had bipolar spindles (Figure 7 and Supplemental Figure 3). Taken together, in addition the previously known role of Cyclin A/Cdk1 activity in destabilizing k-MT attachments in prometaphase (Kabeche and Compton, 2013), our data here indicate that Cyclin A/Cdk1 activity promotes the localization of MPS1, KNL1 and its phosphorylation, and BUB1 to kinetochores to promote chromosome congression and timely mitotic progression.

**Figure 7:**
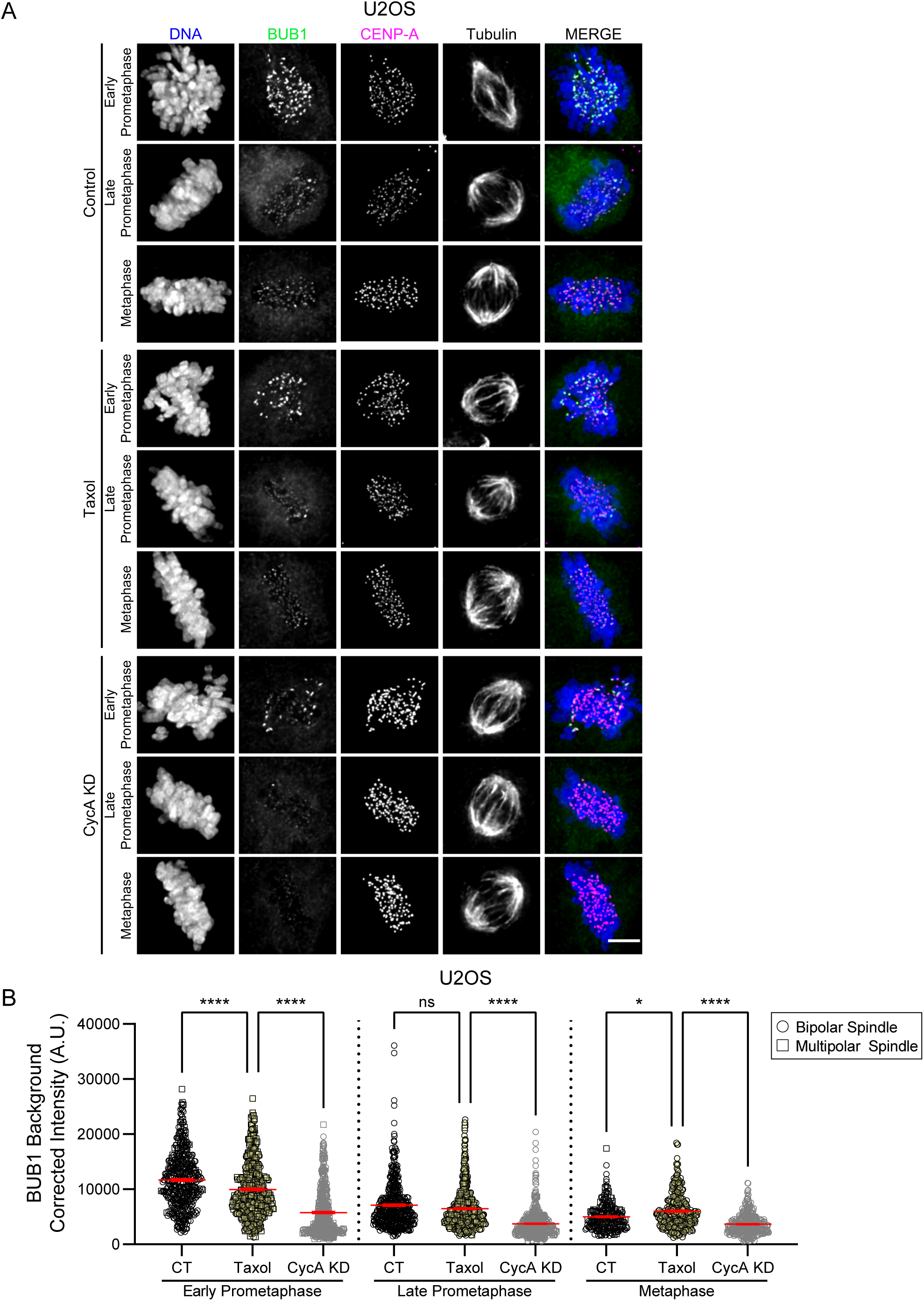
BUB1 localization to kinetochores is independent of changes to k-MT attachment stability. **(A)** Representative images of U2OS cells treated with 0.1% DMSO, 5 nM Taxol for 1 hour, or Cyclin A-depleted (CycA KD). Shown is DNA (blue), BUB1 (green), CENP-A (magenta), and microtubules (greyscale). Scale bar: 5 µm. **(B)** Quantification of BUB1 protein levels at kinetochores of control (CT, black circles or squares), Taxol (yellow circles or squares), or CycA KD (gray circles or squares) U2OS cells. Circles = bipolar spindles and squares = multipolar spindles. n = 225-450 kinetochores from 15-30 cells per condition for each mitotic phase from four independent experiments; ns p > 0.05, *p ≤ 0.05, ****p ≤ 0.0001 using a one-way ANOVA and Šídák’s multiple comparisons test; error bars (red) indicate SEM. See also Supplemental Figure 3.

## DISCUSSION

Mammalian cells enter mitosis with high levels of both Cyclin A/Cdk1 and Cyclin B/Cdk1 activity, and despite the potential for overlap in substrate specificity, each kinase appears to make distinct contributions to mitosis. Cyclin B/Cdk1 activity is required for nuclear envelope breakdown (Hégarat *et al*., 2020) and for maintenance of a mitotic state because cells exit mitosis rapidly if it is inhibited or following Cyclin B destruction by the APC/C once the SAC is satisfied (Brito and Rieder, 2006; Gong and Ferrell, 2010). While Cyclin A/Cdk1 activity is required for mitotic entry (Gong and Ferrell, 2010; Hégarat *et al*., 2020) and facilitates nuclear envelope breakdown and chromatin condensation (Gong and Ferrell, 2010), its activity is not required to maintain a mitotic state (Gong and Ferrell, 2010). Rather, data provided here and elsewhere indicate that its activity during prometaphase, although coincident with its degradation, imparts increased efficiency to events in prometaphase. Here we extend our previous findings that Cyclin A/Cdk1 activity regulates k-MT attachment stability in prometaphase (Kabeche and Compton, 2013) and demonstrate that Cyclin A/Cdk1 activity promotes the formation of bioriented k-MT attachments and chromosome congression. Specifically, we find that cultured cells depleted of Cyclin A display unaligned chromosomes with erroneous syntelic, monotelic, and lateral MT attachments and in some instances, unattached chromosomes consistent with studies performed in *cyclin A2^-/-^* mouse oocytes (Zhang *et al*., 2017). Furthermore, we find that a consequence of impaired bioriented k-MT attachment formation and chromosome congression is a prometaphase delay. Thus, Cyclin A/Cdk1 activity also supports timely mitotic progression (Figure 8). Without Cyclin A/Cdk1 activity regulating these events, there is an increase in mitotic errors (shown here and (Kabeche and Compton, 2013)), including lagging chromosomes in anaphase that can cause chromosome missegregation and aneuploidy (Thompson and Compton, 2011). In this context, Cyclin A/Cdk1 activity isn’t strictly required for mitosis in the same way that Cyclin B/Cdk1 activity is because cells lacking Cyclin A (and by extension Cyclin A/Cdk1 activity) ultimately complete mitosis, but rather it is required for timely progression through mitosis and high fidelity chromosome segregation.

**Figure 8:**
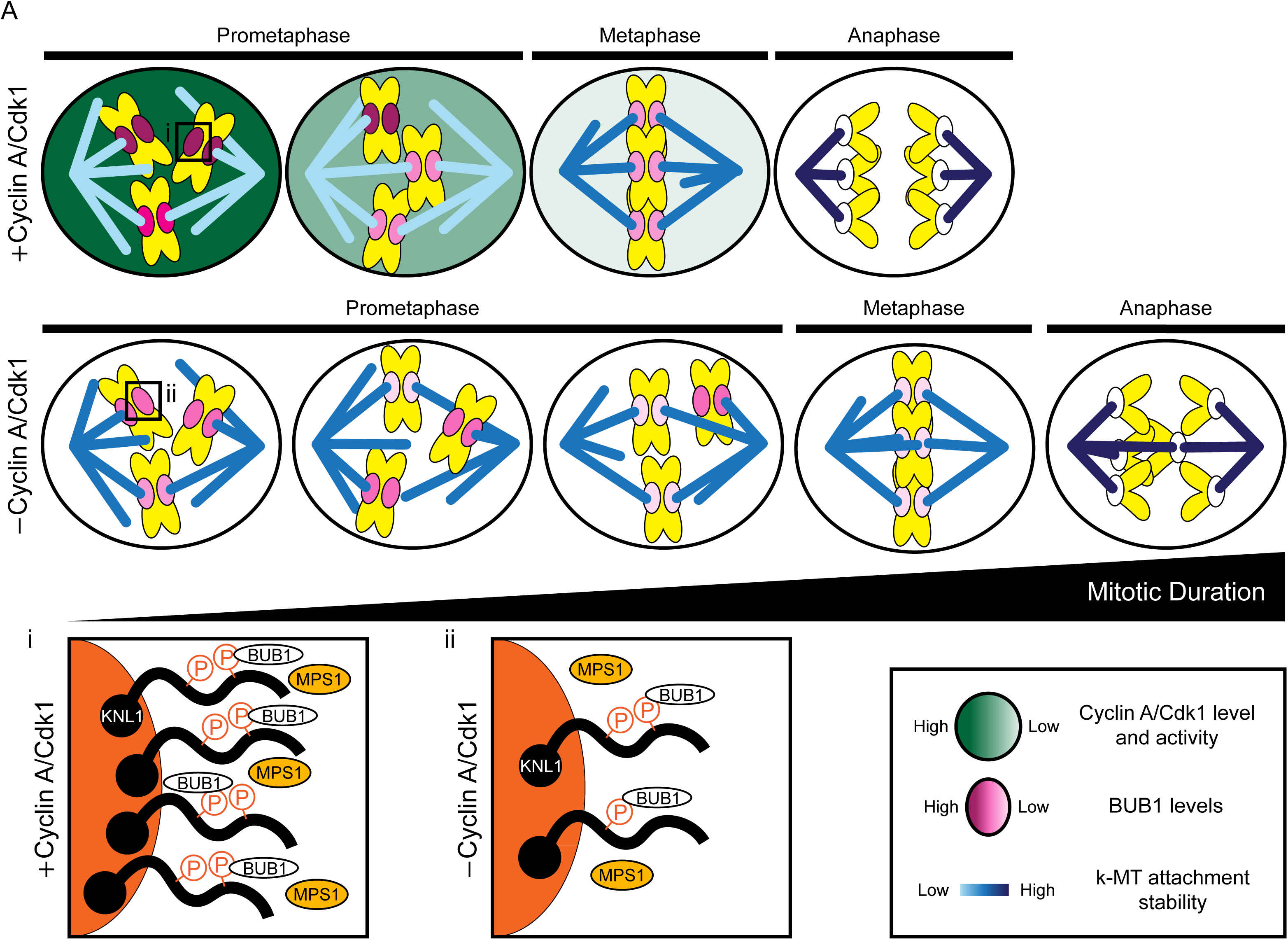
Model for the roles of Cyclin A/Cdk1 in mitosis. **(A)** Cyclin A/Cdk1 activity regulates multiple events in prometaphase, including the establishment of bioriented k-MT attachments, k-MT attachment stability, chromosome alignment, and mitotic progression to ensure high fidelity chromosome segregation. Compared to cells with Cyclin A/Cdk1 activity (top panel), cells lacking Cyclin A/Cdk1 activity (bottom panel) exhibit hyperstable k-MT attachments and delayed prometaphase progression due to impaired formation of bioriented k-MT attachments and chromosome congression to the spindle equator. These defects lead to mitotic errors, including lagging chromosomes in anaphase as depicted in the bottom panel anaphase. Cyclin A/Cdk1 activity in part regulates these prometaphase events by promoting the kinetochore localization of MPS1 kinase, KNL1 and its phosphorylation at MELT sites, and BUB1 kinase (insets i and ii). In the diagram, Cyclin A/Cdk1 activity and levels are represented as shades of green, mitotic duration is indicated by the black triangle, k-MT attachment stability is represented as shades of blue, and BUB1 protein levels are represented by shades of magenta.

Mechanistically, Cyclin A deficiency could impede the establishment of bioriented k-MT attachments during prometaphase through hyper-stable k-MT attachments that prevent error correction (Kabeche and Compton, 2013; Zhang *et al*., 2017). Alternatively, the physical congression of chromosomes from the spindle pole to the spindle equator also facilitates the establishment of bioriented k-MT attachments independent of k-MT attachment dynamics (Risteski *et al*., 2021). Our data demonstrates that Cyclin A-depleted cells fail to recruit a full complement of BUB1 and KNL1 to kinetochores in prometaphase, both of which play a role in chromosome alignment (Johnson *et al*., 2004; Meraldi and Sorger, 2005; Klebig *et al*., 2009; Caldas *et al*., 2013; Zhang *et al*., 2013; Silió *et al*., 2015). Moreover, we demonstrate that BUB1 localization defects in Cyclin A-depleted cells are independent of altering k-MT attachment stability. Thus, Cyclin A/Cdk1 activity regulates both k-MT attachment stability and chromosome congression to achieve full chromosome biorientation. Importantly, the siRNA knockdown strategy utilized here reduced Cyclin A levels in mitotic cells by ∼97-98% although BUB1 levels at kinetochores only decreased by approximately 45%, on average (Supplemental Figure 1A and Figure 3). Thus, Cyclin A/Cdk1 activity merely enhances the recruitment and/or maintenance of BUB1 at kinetochores and Cyclin A-depleted cells still recruit sufficient BUB1 to kinetochores to support SAC function (Zhang *et al*., 2019a). Although, it is possible that Cyclin A depletion causes a weakened SAC response through the resulting decrease in BUB1 localization to kinetochores, in part because partial depletion of BUB1 weakens SAC signaling (Jema *et al*., 2023). Future studies will be required to test the strength of the SAC upon Cyclin A depletion. Perhaps there are two mechanisms for BUB1 recruitment and maintenance at kinetochores; one regulated by Cyclin A/Cdk1 activity for the purpose of promoting bioriented k-MT attachments and chromosome congression, and a second that interfaces with RZZ complex to regulate SAC function (McGory *et al*., 2023).

Cyclin A/Cdk1 may directly or indirectly regulate BUB1 activity and/or localization during prometaphase for these functions. In support of direct regulation, a phosphoproteomic screen identified BUB1 as a potential substrate that is selectively phosphorylated by Cyclin A/Cdk1 during prometaphase (Dumitru *et al*., 2017). Alternatively, we demonstrate that at kinetochores in Cyclin A-depleted mitotic cells MPS1 kinase, KNL1, and phosphorylation of KNL1 MELT motifs are all reduced. Because a complex of BUB1:BUB3 binds phosphorylated KNL1 MELT motifs (Primorac *et al*., 2013) this could explain the decrease in BUB1 at kinetochores upon Cyclin A depletion. In this scenario, Cyclin A/Cdk1 activity would promote BUB1 localization to kinetochores indirectly through a pathway involving MPS1 kinase (Yamagishi *et al*., 2012; Alfonso-Pérez *et al*., 2019; Hayward *et al*., 2019). Regardless of the specific mechanism(s) regulating BUB1 recruitment to kinetochores in prometaphase, these data and previous studies provide insights into how a distributive control system acting to regulate events at all kinetochores coordinately in mitotic cells (Cyclin A/Cdk1 activity) influences the localization and activity of regulatory molecules such as BUB1, MPS1, Cyclin B/Cdk1 (Kucharski *et al*., 2022) and Plk1 (Dumitru *et al*., 2017) that operate chromosome autonomously and locally at kinetochores (Carmena *et al*., 2012; Kabeche and Compton, 2013; Krenn and Musacchio, 2015).

The disruption of bioriented k-MT attachments and chromosome congression in Cyclin A-depleted mitotic cells is also the most likely explanation for the extended duration of prometaphase presumably because there is sufficient ongoing SAC activity to prevent anaphase onset until chromosomes attain bioriented k-MT attachments. Yet, in a recent study partial depletion of BUB1 resulted in the opposite phenotype and an overall shorter mitotic duration (Jema *et al*., 2023). However mitotic duration was measured in combination with CENP-E inhibition preventing a direct comparison with the BUB1 reduction we observe upon Cyclin A depletion in CENP-E proficient cells. Paradoxically, the overexpression of Cyclin A in mammalian cells also prolongs mitotic duration and causes chromosome alignment defects (den Elzen and Pines, 2001). How both depletion and over-expression of Cyclin A cause similar phenotypes is unknown. One possibility is that the effect of Cyclin A/Cdk1 activity on k-MT attachment dynamics explains the similar phenotypes. Cyclin A over-expression increases the detachment of MTs from kinetochores (Kabeche and Compton, 2013) and thus, excess Cyclin A may increase the frequency of unattached kinetochores impairing chromosome congression and delaying mitosis due to SAC signaling. Yet, Cyclin B1/Cdk1 is degraded and the SAC protein Mad2 is not localized to kinetochores in cells with a prolonged mitosis due to Cyclin A overexpression (den Elzen and Pines, 2001) suggesting a more complex scenario where Cyclin A/Cdk1 regulation of mitotic duration may be independent of active SAC signaling. In conclusion, to support timely mitotic progression and prevent genomic instability caused by a prolonged mitosis (Uetake and Sluder, 2010; Meitinger *et al*., 2022) Cyclin A/Cdk1 activity is required in prometaphase, but its timely destruction is also required to prevent ongoing Cyclin A/Cdk1 function throughout prometaphase and potentially during later stages of mitosis.

## MATERIALS AND METHODS

### Cell culture

U2OS (ATCC®, HTB-96) and HeLa cKMKO A6.1 (provided by I. Cheeseman; Massachusetts Institute of Technology, Cambridge, MA) cells were grown in Dulbecco’s modified Eagle’s medium, (DMEM; Corning) supplemented with 10% FCS (HyClone), or DMEM supplemented with 10% tetracycline-free FBS (Denville Scientific), respectively. DMEM was also supplemented with 10mM HEPES (Sigma-Aldrich), 250 μg/L Amphotericin B (Sigma Aldrich), 50 U/mL penicillin, and 50 μg/mL streptomycin (Thermo Fisher Scientific). For live-cell imaging, U2OS-H2B mCherry cell lines were grown in FluoroBrite DMEM (Gibco), supplemented with 10% FCS (HyClone), 10mM HEPES (Sigma-Aldrich), 250 μg/L Amphotericin B (Sigma Aldrich), 50 U/mL penicillin, and 50 μg/mL streptomycin (Thermo Fisher Scientific). Cell lines were validated as mycoplasma free (Sigma-Aldrich, Mycoplasma Kit). Cell lines were cultured in a 37°C-humidified environment with 5% CO_2_.

### Transfections

siRNA transfections were performed by plating cells on 18 mm glass coverslips into 12-well plates with DMEM, or for live-cell imaging, by plating cells onto 35 mm glass-bottom dishes (MatTek) with FluoroBrite DMEM. Twenty-four hours after plating, cells were transfected with 150 nM Cyclin A-CCNA2, ID: s2513 (Silencer Select Validated; 5’-GA UAUACCCUGGAAAGUCUtt-3’) (Thermo Fisher Scientific) with JetPrime (Polyplus) following manufacturer’s siRNA transfection 12-well format protocol for cells on coverslips or 6-well format protocol for cells on glass-bottom dishes. Control transfections were performed following the JetPrime protocol without siRNA. For fixed cell immunofluorescence (IF) experiments, fixation was performed 48 hours after siRNA treatment. For live-cell timelapse imaging, cells on dishes were mounted and imaged on a microscope for 24 through 48 hours after siRNA treatment.

To generate U2OS H2B-mCherry cell line, U2OS cells were transfected with H2B-mCherry plasmid, a gift from Robert Benezra (Addgene plasmid # 20972), using Fugene (Promega) in OptiMEM (Thermo Fisher Scientific). After transfection and growth, cells were cultured with G418 at 1mg/mL (InvivoGen) and colonies were selected.

### Drug Treatment

Paclitaxel (Taxol) was used at 5 nM (BioTang) for 1 hour prior to fixation.

### Antibodies

Primary antibodies for immunofluorescence are as follows: ACA (1:750; Geisel School of Medicine), BUB1 (1:1000; Abcam; ab9000), BUB1 (1:250; Abcam; ab54893), CENP-A (1:500; Thermo Fisher Scientific; MA1-20832); Cyclin A (1:400; GeneTex; GTX634420), H2ApT120 (1:1000; Active Motif; #61195), KNL1 (1:250; Abcam; ab70537), pKNL1 (pMELT pT943/pT1155; 1:400; Cell Signaling; #40758), MPS1 (1:50; EMD-Millipore; 05-683), Tyrosinated Tubulin (Tubulin; 1:1000; Novus; NB600-506). Secondary antibodies for immunofluorescence are Alexa Fluor 488, 594, or 647 antibodies, raised in donkey or goat, against mouse, rabbit, or human (1:2000; Thermo Fisher Scientific). DAPI was used to stain DNA (500 ng/mL; Sigma Aldrich; D9542).

Primary antibodies for Western blotting are as follows: BUB1 (1:2000; Abcam; ab9000) Cyclin A2 (1:2500; Abcam; ab9000), Cyclin B1 (1:100; Santa Cruz; sc-245), GAPDH (1:1000; Santa Cruz; sc-365062), H3pS10 (1:1000; Cell Signaling: #53348). Secondary antibodies for Western blotting are IRDye 680RD Goat anti-Mouse (1:10000 or 1:20000; #926-68070; LI-COR, IRDye 800CW Goat anti-rabbit (1:10000; #926-32211, LI-COR), and IRDye 800CW Goat anti-mouse (1:2500 or 1:20000; #926-32210; LI-COR).

### Immunofluorescence

For immunofluorescence staining of BUB1, Cyclin A2, KNL1 pMELT, H2ApT120, ACA, and CENP-A, cells were fixed for 10 minutes with room temperature (RT) 3.5% PFA (paraformaldehyde, pH 6.7; Alfa Aesar). Cells were then washed 3 x with Tris buffered saline (TBS) with 1% bovine serum albumin (BSA; VWR) and 0.5% Triton X-100 (Sigma) for 5 minutes. Cells were incubated in TBS-BSA for 5 minutes, followed by 1 x 5-minute wash with TBS-BSA + 0.1% Triton X-100. Washes were followed by a 2-hour RT incubation with primary antibodies diluted in TBS-BSA + 0.1% Triton X-100. Cells were washed 3 x with TBS-BSA + 0.1% Triton X-100 for 10 minutes and then incubated for 1 hour with secondary antibodies and DAPI diluted in TBS-BSA + 0.1% Triton X-100 while protected from light. Next, cells were washed 3 x with TBS-BSA+ 0.1% Triton X-100 for 10 minutes and rinsed with TBS-BSA. Coverslips were then mounted on slides using Prolong Gold Antifade Mountant (Thermo Fisher Scientific).

For immunofluorescence staining of experiments using the total KNL1 antibody, cells were fixed and permeabilized simultaneously in PTEM-F buffer (50 mM PIPES, pH 6.8; 0.2% Triton X-100; 10 mM EGTA; 1 mM MgCl^2^; and 3.7% formaldehyde) for 10 minutes at RT. Cells were then washed 2 x TBS-BSA + 0.5% Triton X-100 for 10 minutes and rinsed with TBS-BSA. Washes, primary and secondary incubations, and coverslip mounting were carried out as above.

For immunofluorescence staining of experiments using the MPS1 antibody, cells were fixed and permeabilized simultaneously in PTEM-F buffer for 10 minutes at RT. Cells were then washed twice with 1X PBS and then incubated with TrueBlack IF Background Suppressor (Biotium) for 30 minutes at RT. Subsequently, cells were incubated with primary antibodies diluted in TrueBlack® IF Blocking Buffer (Biotium) for 2 hours at RT. Primary incubation was followed by two 1X PBS rinses and three 1X PBS washes for 5 minutes. Cells were then incubated with secondary antibodies diluted in TrueBlack® IF Blocking Buffer (Biotium) for 1 hour at RT. Secondary antibody and DAPI incubation was followed by two 1X PBS rinses and three 1X PBS washes for 5 minutes. Coverslips were then mounted on slides using Prolong Gold Antifade Mountant (Thermo Fisher Scientific).

### Calcium-Stable Microtubule Assay

For visualization of calcium stable microtubules, cells were extracted using a calcium buffer (0.1mM CaCl_2_ [U2OS] or 0.05mM CaCl_2_ [HeLa] with 100 mM PIPES, 1 mM MgCl2, and 0.5% Triton X-100 at pH 6.8) for ∼5 minutes at RT, followed by fixation using a 1% glutaraldehyde solution for 10 minutes. Fixation was followed by two washes with 0.5 mg/mL NaBH_4_ for 10 minutes, one 5 minute wash in TBS-BSA and two 5-minute washes with TBS-BSA + 0.5% Triton X-100. Washes, primary and secondary incubations, and coverslip mounting were carried out as above, except for a 3-hour RT incubation with primary antibodies.

### Western blotting

After transfections, cells were collected, lysed, and boiled in SDS sample buffer (1M Tris, 50% glycerol, 10% SDS, 0.5% bromophenol blue, β-mercaptoethanol). Samples were loaded onto SDS-PAGE gels and run. Proteins were wet transferred onto nitrocellulose membranes overnight (BioRad). Membranes were dried overnight and rehydrated in 1X TBS for 5 minutes. The membranes were subsequently blocked in Odyssey Blocking Buffer (LI-COR) for 1 hour at RT. Following blocking, membranes were incubated for 1-2 hours at RT with primary antibodies diluted in Odyssey Blocking Buffer + 0.2% Tween (VWR). Next, membranes were washed 4 x with TBS-T (0.1% Tween 20) for 5 minutes. Membranes were then incubated with secondary antibodies diluted in Odyssey Blocking Buffer + 0.2% Tween 20 at RT and protected from light. Following incubation with secondary antibodies, membranes were then washed 4 x with TBS-T for 5 minutes and rinsed with TBS. Membranes were imaged on an Odyssey CLx Digital Imager (LI-COR). Quantification of band signal intensities were performed using Image Studio Lite software (LI-COR).

### Microscopy and Quantification

Timelapse live-cell fluorescence imaging was performed using a Nikon Ti2-E Inverted Microscope using a Plan APO λD 60x, 1.42 NA, oil immersion objective (Nikon). The microscope was equipped with a Prime BSI (Teledyne Photometrics) camera, a SoRa CSU-W1 LUN-F XL spinning disk confocal system (Yokogawa; Nikon) with an iChrome MLE laser engine (Toptica). Cells on glass bottom dishes were kept in a humidified stagetop incubator and maintained at 37°C with 5% CO_2_ throughout imaging (Tokai Hit). Images of 2 stage positions using 1 µm steps for a total of 14 optical slices, were acquired at 2 minute intervals for 24 hours. Mitotic duration and anaphase errors were quantified by manually tracking cells using Elements software (Nikon). Images on panels are maximum projections from z-stacks of all 14 optical slices and contrasted to visualize H2B-mCherry fluorescence.

Images for calcium stable microtubules were acquired using a Nikon Eclipse Ti inverted microscope using a Plan Apo VC 100x, 1.4 NA, oil immersion objective (Nikon). The microscope was equipped with an Evolve Delta camera (Photometrics) and a QuorumWave FX-X1 spinning disk confocal system with an ILE laser engine (Andor Technology). More than 90 steps of .1 µm were used to capture entire spindles and visualize attachments. Figure images are deconvolved (Nikon Elements) maximum intensity projections from z-stacks of 2-10 imaging planes. Figure panels from different cells have different contrast enhancements (Fiji) to better visualize attachments.

All other images were acquired using a Nikon Eclipse Ti Inverted microscope using a Plan Apo VC 60x, 1.4 NA, oil immersion objective (Nikon) and equipped with an ORCA-Fusion Gen III sCMOS camera (Hamamatsu) and a SOLA LED light engine (Lumencor). Images were acquired using 0.25 µm steps. For quantification of intercentromere distance, in Fiji (Schindelin *et al*., 2012), a circular region, 8 pixels in diameter was manually drawn around CENP-A foci of sister centromeres to calculate distance. For quantification of proteins, except KNL1 and MPS1, measurements were taken using the CRaQ Fiji plugin as described in (Bodor *et al*., 2012; Schindelin *et al*., 2012). In brief, background corrected intensity measurements are calculated as the difference between maximum and minimum intensity values at small areas bounding kinetochores (using the CENP-A or BUB1 signal to define kinetochore regions) from maximum intensity projections of z-stacks. For quantification of KNL1 and MPS1, background corrected intensity measurement values are calculated as the difference between intensity measurements of KNL1 at a circular kinetochore region 7 pixels in diameter (defined by the BUB1 signal in a separate channel) and a circular region of 8 pixels in diameter around the kinetochore from single z-slices as described previously, using Fiji (Hoffman *et al*., 2001). Figure images are deconvolved (Nikon Elements) maximum intensity projections from z-stacks of 20-25 imaging planes. Figure panels from different cell lines have different contrast enhancements (Fiji), but contrast within a cell line is consistent for all conditions/mitotic stages, except for the tubulin channel in Taxol-treatment experiments.

### Statistical Analysis

All experiments are data from at least three independent biological replicates, unless otherwise specified in the figure legend. Sample sizes are stated in figure legends. All statistical analyses were performed using GraphPad 9. Statistical tests used were Fisher’s Exact test, t-test, or ANOVA, with multiple comparisons corrections for specified pairs of conditions analyzed. Error bars on graphs represent mean ± SEM. p-values are shown as: ns p > 0.5; *p ≤ 0.05, **p ≤ 0.01,***p ≤ 0.001, and ****p ≤ 0.0001 in the figure legends.

## ACKNOWLEDGMENTS

We thank the members of the Compton and Godek labs for helpful discussions and technical advice. The authors acknowledge the Shared Resources facilities at the Dartmouth Cancer Center with NCI Cancer Center Support Grant P30 CA023108, and the Life Sciences Light Microscopy Facility at Dartmouth, particularly Ann Lavanway. This work was supported through NIH awards R37GM051542 to D.A.C., R01HD101436 to K.M.G, and S10OD032310 to Dr. Y. Ahmed for purchase of the super-resolution spinning disk confocal, bioMT through NIH grant P20-GM113132, DartCF through NIH grant P30-DK117469.

## FIGURE LEGENDS

**Supplemental Figure 1:**
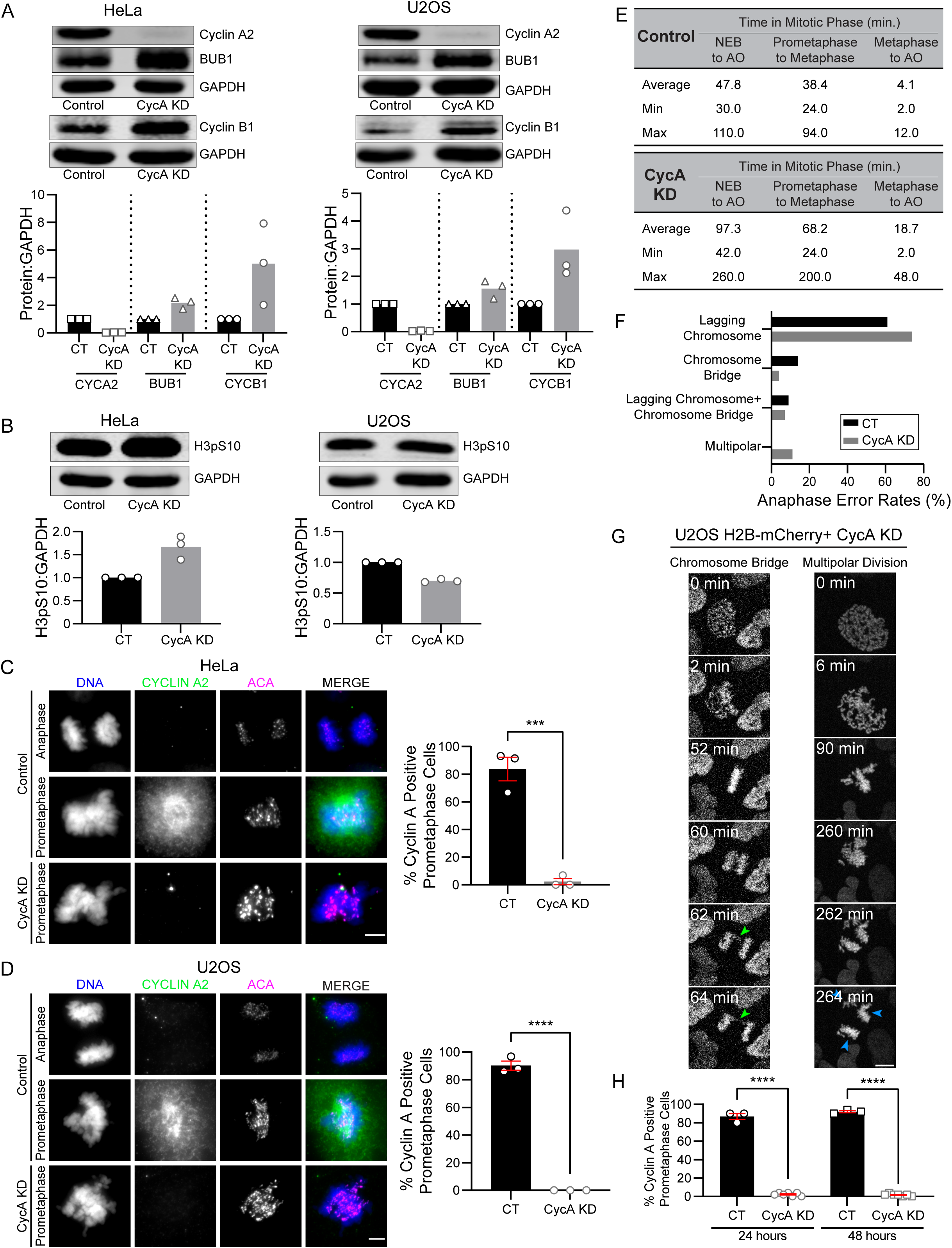
Validation of Cyclin A siRNA depletion in HeLa and U2OS cells and anaphase error rates from timelapse live-cell imaging of Cyclin A depleted U2OS cells. Related to Figure 1. **(A)** Representative Western blots from asynchronous HeLa (left) or U2OS (right) populations for Cyclin A2, BUB1, GAPDH (loading control), or Cyclin B1 protein levels in control (CT) or Cyclin A depleted (CycA KD) cells treated with 150 nM Cyclin A siRNA for 48h. Bar graphs below show quantification of Cyclin A2, BUB1, and Cyclin B1 protein levels relative to GAPDH for CT (black bars) or CycA KD cells (gray bars). **(B)** Representative Western blots from asynchronous HeLa (left) or U2OS (right) populations for H3pS10 and GAPDH (loading control) protein levels in CT or CycA KD cells treated with 150 nM Cyclin A siRNA for 48h. Bar graphs below show quantification of H3pS10 protein levels relative to the GAPDH for CT cells (black bars) or CycA KD cells (gray bars). **(C, D)** Representative images of CT and CycA KD HeLa cells **(C)** or U2OS cells **(D)**. Shown is DNA (blue), Cyclin A2 (green), and ACA (magenta; centromere marker) in prometaphase or anaphase. Bar graphs (right) show quantification of prometaphase cells scored for Cyclin A2 in CT (black bars) and CycA KD (gray bars) HeLa cells **(C)** or U2OS cells **(D)**. n = 100 prometaphase cells scored per condition. Scale bars: 5 µm. **(E)** Summary of average, minimum, and maximum prometaphase, metaphase, and total mitotic duration of control and CycA KD U2OS H2B-mCherry cells from timelapse live-cell imaging. Data from Figure 1C. **(F)** Percentage of anaphase error rates, including lagging chromosomes, chromosome bridges, lagging chromosomes + chromosome bridges, and multipolar anaphases in CT and CycA KD U2OS H2B-mCherry cells from timelapse live-cell imaging. Data from Figure 1C. **(G)** Selected panels from timelapse live-cell fluorescence imaging of U2OS H2B-mCherry CycA KD cells showing a chromosome bridge (left; green arrowheads) or a multipolar anaphase (right; blue arrowheads). Scale bar: 10 µm. **(H)** In parallel samples, the percent of prometaphase CT (black bars) or CycA KD (gray bars) cells expressing Cyclin A was quantified by immunofluorescence at 24 hours (imaging initiation) and 48 hours (imaging termination). For CT n = 150 cells (24 hours) and n = 150 cells (48 hours). For CycA KD n = 350 cells (24 hours) and n = 351 cells (48 hours). Data from **(A, B, C,** and **D)** is from three independent experiments; data for **(E, F,** and **H)** is from the three (CT) and five (CycA KD) independent experiments in Figure 1C; ***p ≤ 0.001 or ****p ≤ 0.0001 using a two-tailed t test; error bars (red) indicate SEM.

**Supplemental Figure 2:**
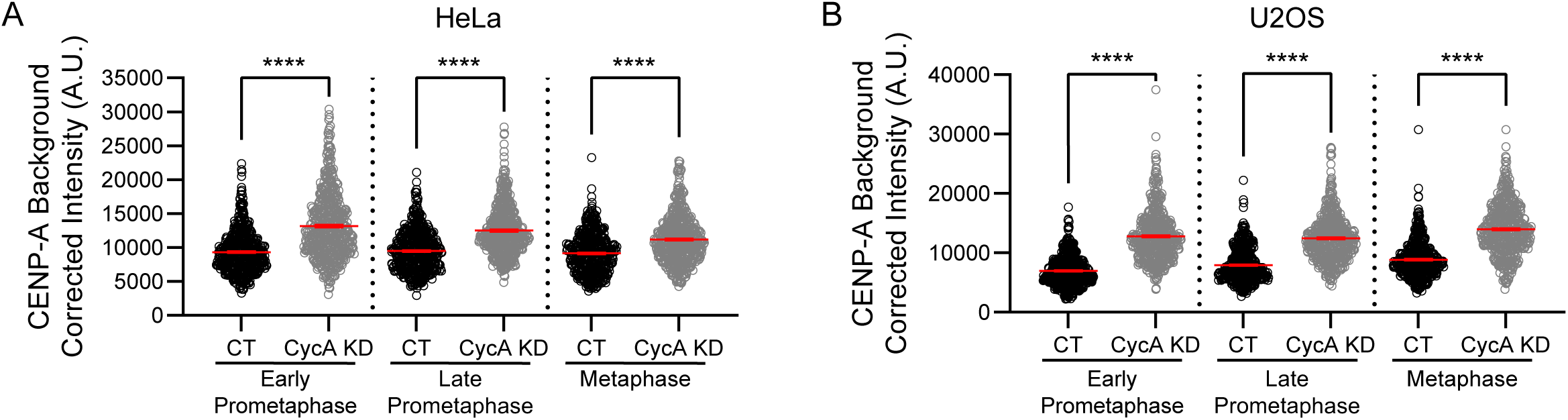
CENP-A is recruited to kinetochores in the absence of Cyclin A. Related to Figure 3. **(A, B)** Quantification of CENP-A protein levels at kinetochores of control (CT, black circles) and Cyclin A depleted (CycA KD, gray circles) HeLa cells **(A)** or U2OS cells **(B)**. n > 500 kinetochores from 29-33 HeLa cells per mitotic phase. n = 500 kinetochores from 30-32 U2OS cells per mitotic phase. Data for **(A** and **B)** is from the three independent experiments in Figure 3B and 3F; ns p > 0.05, **p ≤ 0.01, and ****p ≤ 0.0001 using a one-way ANOVA and Šídák’s multiple comparisons test; error bars (red) indicate SEM.

**Supplemental Figure 3:**
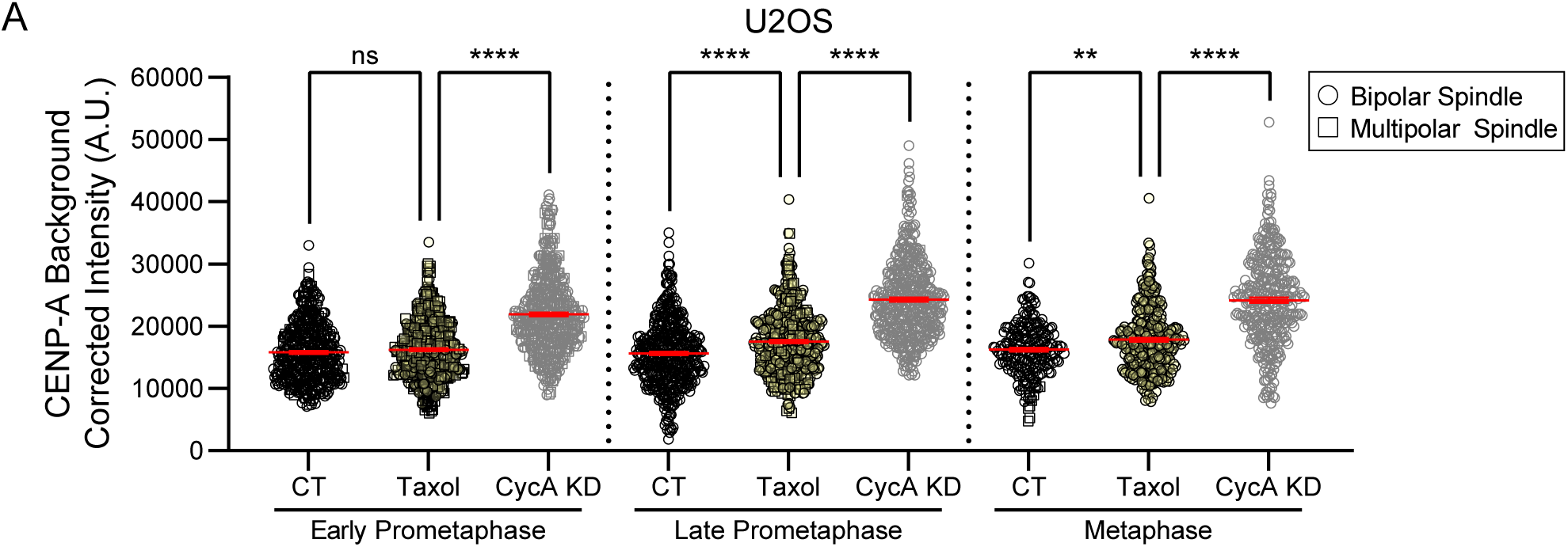
CENP-A recruitment to kinetochores is independent of changes to k-MT attachment stability. Related to Figure 7. **(A)** Quantification of CENP-A protein levels at kinetochores of U2OS cells treated with 0.1% DMSO (CT, black circles or squares), 5nM Taxol for 1 hour (yellow circles or squares), or CycA KD (gray circles or squares). Circles = bipolar spindles and squares = multipolar spindles. n = 225-450 kinetochores from 15-30 cells per condition for each mitotic phase. Data is from the four independent experiments in Figure 7B; ns p > 0.05, **p ≤ 0.01, and ****p ≤ 0.0001 using a one-way ANOVA and Šídák’s multiple comparisons test; error bars (red) indicate SEM.

## Notes

### Competing Interest Statement

The authors have declared no competing interest.

